# Flexibility of learning in complex worlds

**DOI:** 10.1101/2023.06.12.544544

**Authors:** Olof Leimar, Andrés E Quiñones, Redouan Bshary

## Abstract

Learning to adjust to changing environments is an important aspect of behavioral flexibility. Here we investigate the possible advantages of flexible learning rates in volatile environments, using learning simulations. We compare two established learning mechanisms, one with fixed learning rates and one with flexible rates that adjust to volatility. We study three types of ecological and experimental volatility: transitions from a simpler to a more complex foraging environment, reversal learning, and learning set formation. For transitions to a complex world, we use developing cleaner fish as an example, having more types of client fish to choose between as they become adult. There are other similar transitions in nature, such as migrating to a new and different habitat. Performance in reversal learning and in learning set formation are commonly used experimental measures of behavioral flexibility. Concerning transitions to a complex world, we show that both fixed and flexible learning rates perform well, losing only a small proportion of available rewards in the period after a transition, but flexible rates perform better than fixed. For reversal learning, flexible rates improve the performance with each successive reversal, because of increasing learning rates, but this does not happen for fixed rates. For learning set formation, we find no improvement in performance with successive shifts to new stimuli to discriminate for either flexible or fixed learning rates. Flexible learning rates might thus explain increasing performance in reversal learning, but not in learning set formation. We discuss our results in relation to current ideas about behavioral flexibility.

## Introduction

The ability of animals to adjust to new and complex environments through learning is an important aspect of adaptive behavioral flexibility (Fawcett et al. 2013). In animal psychology and behavioral ecology, different meanings have been given to the term behavioral flexibility (Audet and Lefebvre 2017; Lea et al. 2020; Uddin 2021), but here we are concerned with the ability to adjust to environmental change using learning, for instance learning to select suitable food items. The question we ask is how well different learning rules (Fawcett et al. 2013), in the sense of different mechanisms of reinforcement learning, with either constant or variable learning rates, serve to adapt behavior in a volatile environment. Specifically, we investigate how big the advantage of having flexible learning rates might be in a volatile environment.

It is known from neuroscience studies that humans and other animals adjust learning rates to the volatility of rewards (e.g., Behrens et al. 2007; Diederen and Schultz 2015; Grossman et al. 2022). An experimental example of volatility in rewards is reversal learning, where an individual first learns to discriminate between a rewarded and a nonrewarded option, and then the rewards are reversed, sometimes with successive episodes of reversal. Performance in such reversal learning is one measure that has been used to describe behavioral flexibility (Deaner et al. 2006; Bond et al. 2007; Izquierdo et al. 2017; Liu et al. 2016; Buechel et al. 2018; Boussard et al. 2021; Triki et al. 2022; Vardi and Berger-Tal 2022), and this performance might be improved by flexible learning rates. Reversal in rewards is a particular form of volatility, but it is similar to forms of volatility that may occur in nature (Raine and Chittka 2012; Cauchoix et al. 2017). The performance in other measures of behavioral flexibility, such as learning set formation (set-shifting), where an individual encounters a sequence of novel discrimination tasks (Harlow 1949; Wilson et al. 1985; Audet and Lefebvre 2017; Bailey et al. 2007), could conceivably also be enhanced by flexible learning rates.

To investigate the potential advantages of variable learning rates, we use a classical learning mechanism with fixed rates as a baseline. Rescorla and Wagner (1972) introduced a model for classical conditioning with complex, multi-dimensional stimuli. The Rescorla-Wagner model can be extended to operant conditioning and is among the most investigated approaches to learning. The model updates estimates of the value of each component or dimension of a complex stimulus. Learning rates can differ between dimensions, but for a given dimension the rate is constant over time. While being a strong candidate for adaptive learning, it is known that Rescorla-Wagner is not optimal in volatile environments (Dayan et al. 2000; Trimmer et al. 2012). Several alternatives to Rescorla-Wagner learning have been proposed, typically involving flexible learning rates. The basic idea is that high learning rates should be advantageous in volatile environments, where there is a need to learn to respond to changes, whereas low learning rates might be advantageous in stationary environments, in particular in stationary environments with stochasticity. In our comparisons here, we use a learning algorithm called Autostep (Mahmood et al. 2012), because of its robustness in adapting learning rates without the need for extensive tuning of parameters. It is a refinement of the so-called delta-bar-delta algorithm (Jacobs 1988; Sutton 1992a, 2022), and it falls into the category of meta-learning approaches (Sutton 2022), i.e. learning to learn.

In the following, we outline the learning models and simulations we use, and then present results from different situations where flexible learning might be advantageous, comparing Rescorla-Wagner with Autostep. The first cases we investigate are inspired by the situation for developing cleaner fish, as they become adult and transition from a simpler to a more complex set of client fish species to choose between and clean (Triki et al. 2019). More generally, these cases illustrate challenges encountered by many learning animals. Examples include migrants experiencing a shift to a new and different foraging environment (Bairlein and Simons 1995; Pierce and McWilliams 2005), and seasonal changes that expose a forager to new food types (Janmaat et al. 2016). We also give an illustration of the performance of Rescorla-Wagner and Autostep when the degree of stochasticity of the environment is high. Although the impacts of stochasticity vs. volatility on learning have not been emphasized in experimental psychology or behavioral ecology, this has been dealt with in neuroscience (Nassar et al. 2010; Piray and Daw 2021). The general conclusion is that stochasticity should favor lower learning rates, allowing a learner to average over more trials. Finally, we compare the performance of Rescorla-Wagner learning to Autostep for a case of reversal learning, extended over several reversals, and we make a similar comparison for a case of learning set formation.

We discuss our results in relation to existing ideas about the significance of flexible learning, making the point that relatively simple mechanisms of adjustment of learning rates could, wholly or partially, explain some of the observed phenomena of flexibility of learning. Such adjustments could represent specific adaptations to environmental volatility, or they could be consequences of broader cognitive adaptations, for instance relating to attention and memory. We also comment on the Autostep model in relation to previous and current ideas in experimental psychology (e.g., Mackintosh 1975; Pearce and Hall 1980; Pearce and Mackintosh 2010; Holland and Schiffino 2016; Soltani and Izquierdo 2019). Finally, we argue that performance in the face of environmental volatility, including transitions to new and complex environments, is a good candidate for the selective advantage of flexible learning.

## Learning models and approaches

The kind of learning we study is where an individual learns the values of stimuli that can be distinguished by certain characteristics, which we refer to as features of compound (multi-dimensional) stimuli. The characteristics define different stimulus dimensions, which can be things like the color, texture, or shape of potential food items encountered by a forager. For the cleaner fish example, a compound stimulus would be a client fish. The individual cleaner fish learns an estimate of a value for each feature of compound stimuli, and uses the sum of these values to estimate the value of the client fish. We refer to this as ‘feature learning’. There are alternative ways that individuals might estimate the values of compound stimuli, for instance to form an entirely separate estimate for each type of stimulus (sometimes called ‘object learning’, Farashahi et al. (2020)).

Here we focus on feature learning. The approach corresponds to long-standing ideas about classical conditioning in experimental psychology, when animals respond to the component stimuli that are present in a learning trial. The case most frequently studied is that of absence/presence features (0/1 stimulus components), where a feature has only two states, either being absent or present in the compound, and we make use of this in our learning simulations. There are of course other cases, for instance quantitative stimulus dimensions, and we include overall stimulus size as one such dimension.

Perhaps the most influential formulation of these ideas is the learning mechanism proposed by Rescorla and Wagner (1972). In their approach, if *w_m_* is an individual’s current estimate of the value of a certain absence/presence feature from stimulus dimension *m*, and the feature is present in a learning trial, the individual updates its estimate to *w^t^*, where

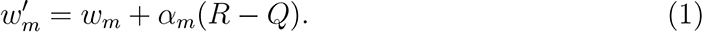

Here *R* is the reward perceived by the individual from interacting with the compound, and *Q* is the individual’s previous estimate of the value of the compound. The quantity

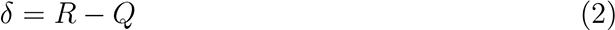

is referred to as the prediction error, and is the difference between the reward *R* currently experienced by the individual and its prior estimate *Q* of the reward. The change in the estimated feature value *w_m_* in equation (1) is thus the learning rate *α_m_* times the prediction error, and tends to move the estimate towards the true value. Learning rates could differ between stimulus dimensions and could also change over time. The main question we ask is how big the advantage of flexible learning rates might be.

If *x_m_*indicates the feature from stimulus dimension *m*, so that *x_m_* is a 0/1 variable, the estimate of the value of the compound is given by the sum of all feature values that are present:

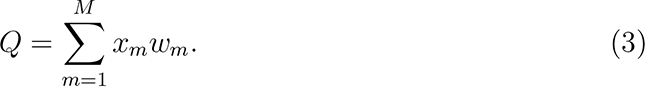

This formula also applies to quantitative stimulus dimensions, for which *w_m_* is the estimated reward per unit of the dimension. For simplicity, we limit ourselves to additive reward structures, although there are other cases that occur in nature, such as when features interact in indicating the value. There is also random variation in rewards. For instance, for client fish visiting cleaners, there is work showing that the number of parasites that cleaners remove and feed on is correlated with client size, but the correlations are not extremely high (Grutter 1994, 1995).

When an individual can choose between two compound stimuli with estimated values of *Q*_1_ and *Q*_2_, we assume that the individual chooses stimulus 1 with probability

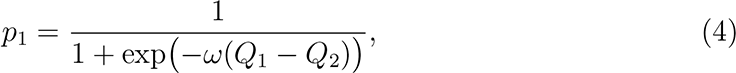

where *ω* is a parameter (we used *ω* = 5 in our simulations; see the curve in Figure 1b below). This is referred to as a soft-max rule, going from estimated values to a choice, and it is commonly used in reinforcement learning models (e.g., Sutton and Barto 2018).

**Figure 1:**
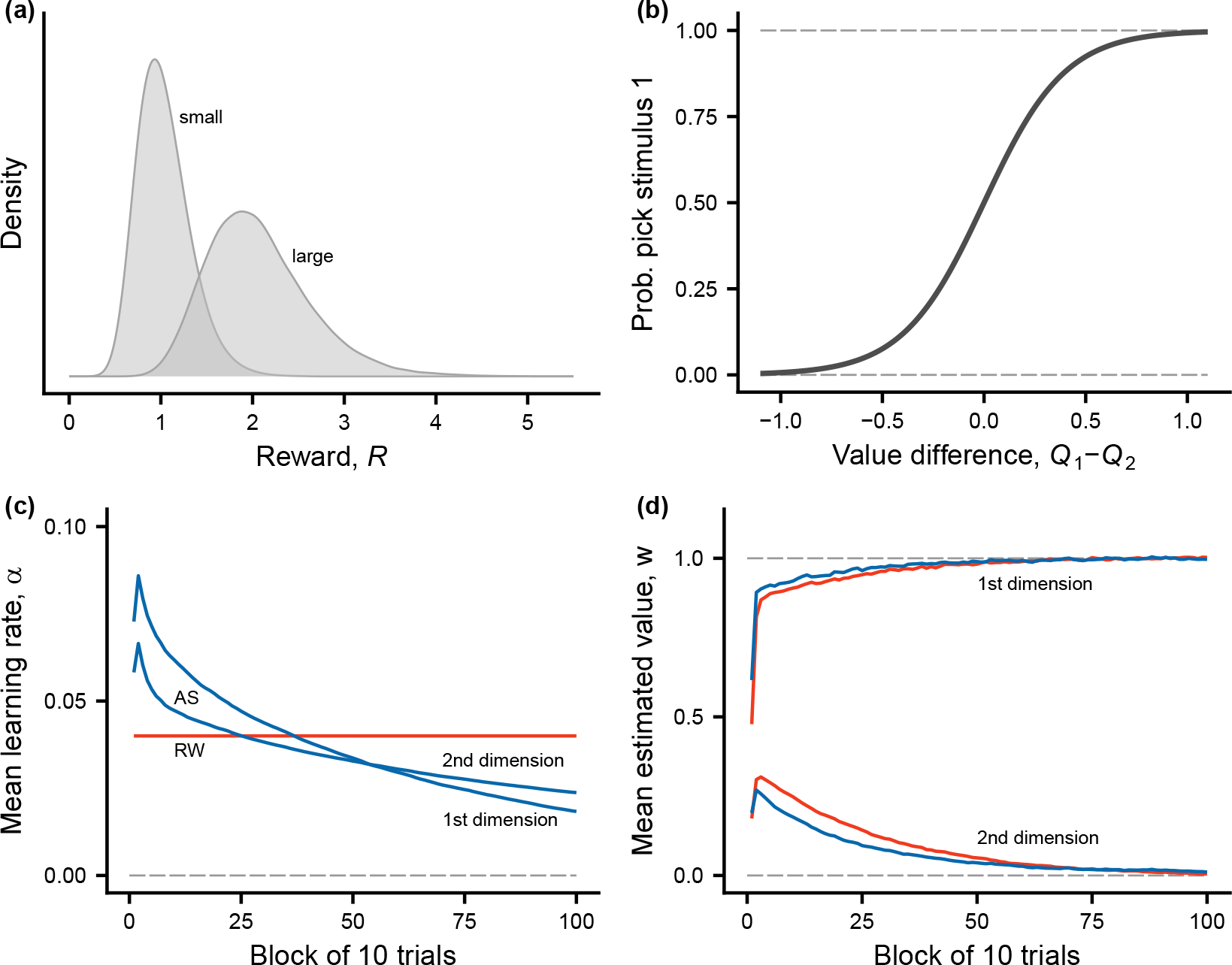
Overview of the first phase, where individuals learn to discriminate between two types of compound stimuli (‘small’ and ‘large’ clients). **(a)** Distribution of rewards from the two types of compound stimuli. **(b)** The function from equation (4), giving the probability of choice from the difference in estimated values of the two compound stimuli present in a trial. **(c)** Learning rates for Rescorla-Wagner (RW) and Autostep (AS) for the two stimulus dimensions. **(d)** Estimated values for Rescorla-Wagner and Autostep for the two stimulus dimensions (first dimension has true value 1.0 and second has true value 0). There are 10 trials in a block and data are averages over 100 replicate learning simulations.

Rescorla and Wagner (1972) assumed that learning rates stay constant over time, but there are a number of suggestions for how they might vary. One idea is that an increase in prediction errors could indicate to an individual that it should change its learning rates. We will investigate how much better an individual with flexible learning rates is at selecting higher value compounds, compared to a Rescorla-Wagner learner. The Autostep method (Mahmood et al. 2012) is a meta-learning approach (Sutton 2022) that adjusts learning rates based on the recent prediction-error history. An overall idea of such meta-learning algorithms is to adjust learning rates in a way that minimizes prediction errors. Autostep is a further refinement of the incremental deltabar-delta (IDBD) method (Sutton 1992a), making it more robust. The intuition behind IDBD is to increase a learning rate *α_m_* if in recent trials the estimate *w_m_* has been increasing (decreasing) and the current prediction error indicates an additional increase (decrease) in *w_m_*, corresponding to a positive correlation between the recent and current changes to *w_m_*. Similarly, the learning rate is decreased for a negative correlation, because this indicates that the current change in *w_m_* overshoots the true value. The algorithm also changes learning rates on the log-scale, which allows for a fairly large range of values for the rates. These properties of IDBD also hold for Autostep. More details on the learning algorithms we use appears in the supplements.

### Learning simulations

As mentioned, the first learning environments we use in our simulations are inspired by the situation for developing cleaner fish, as they become adult and transition from a simpler to a more complex set of client fish species to choose between and clean. Here we describe the simulations for such cases, where there is a transition from a simpler to a more complex learning environment.

#### Stimulus dimensions and compound stimuli

In order to characterize many (up to 10) different compound stimuli, there are 10 stimulus dimensions. The first four dimensions are as follows, together with their true values.

1. The first dimension, *x*_1_, is quantitative, like client size, and has a positive true value, *W*_1_ = 1.0.
2. The second dimension is 0/1, and has a zero true value, *W*_2_ = 0, so it is an uninformative dimension (irrelevant for reward).
3. The third dimension is 0/1 and has a positive true value, *W*_3_ = 1.0, so it is an informative dimension (relevant for reward).
4. The fourth dimension is 0/1 and has a negative true value, *W*_4_ = *−*1.0, so it is also a relevant dimension.

An additional six 0/1 dimensions are described in Table 1 below. From combinations of the four first dimension we have four types of compound stimuli. These could correspond to four client species. Relevant criteria for variation in client value would be size, parasite load, mucus quality/quantity and maneuverability (Roche et al. 2021). Of these, only size is directly visible to cleaners. Correlations between size and the other variables may exist but are weak enough that paying attention to features/dimensions other than size may help cleaners to improve their choices. We hence illustrate the scenario with four species/compound stimuli by considering size as a continuous variable, and colorfulness, swimming with pectoral fin, and a continuous second part of the dorsal fin as dichotomous (0/1).

1. The first type has small size, *x*_1_ = exp(*y*_1_ + *z_x_*), with *z_x_* normally distributed with mean zero and standard deviation *σ_x_* = 0.25, and *y*_1_ so that *x̄*_1_ = *x̄*_small_ = 1 (which happens for *y*_1_ = *−σ*^2^*/*2), and absence of features in the other dimensions. This could be a species of small clients, like less colorful damselfish.
2. The second type has large size, *x*_1_ = exp(*y*_2_ + *z_x_*), again with *z_x_*normally distributed with mean zero and standard deviation *σ_x_*, and *y*_2_ such that *x̄*_1_ = *x̄*_large_ = 2, and presence of a feature in the second dimension, and absence of features in the other dimensions. This could be a species of large clients that are characterized by a feature *x*_2_ that is irrelevant for reward (size is sufficient to predict reward). An example could be a bream, for which being colorful contains no information about parasite load.
3. The third type of compound stimulus is the same as the second for the first two dimensions, but it has a feature present in the third dimension and no feature in the fourth. This is then a species of more valuable large clients. An example could be a thicklip wrasse, which is large, colorful and swims with pectoral fins. These fish have particularly high parasite loads (Grutter 1995).
4. The fourth type of compound stimulus is the same as the second for the first two dimensions, and it has no feature in the third dimension but a feature present in the fourth. This is then a species of less valuable large clients. A snapper could be an example, being large, colorful, not swimming with pectoral fins, and having a continuous second part of the dorsal fin. These fish have less preferred mucus (Grutter and Bshary 2004).

**Table 1:**
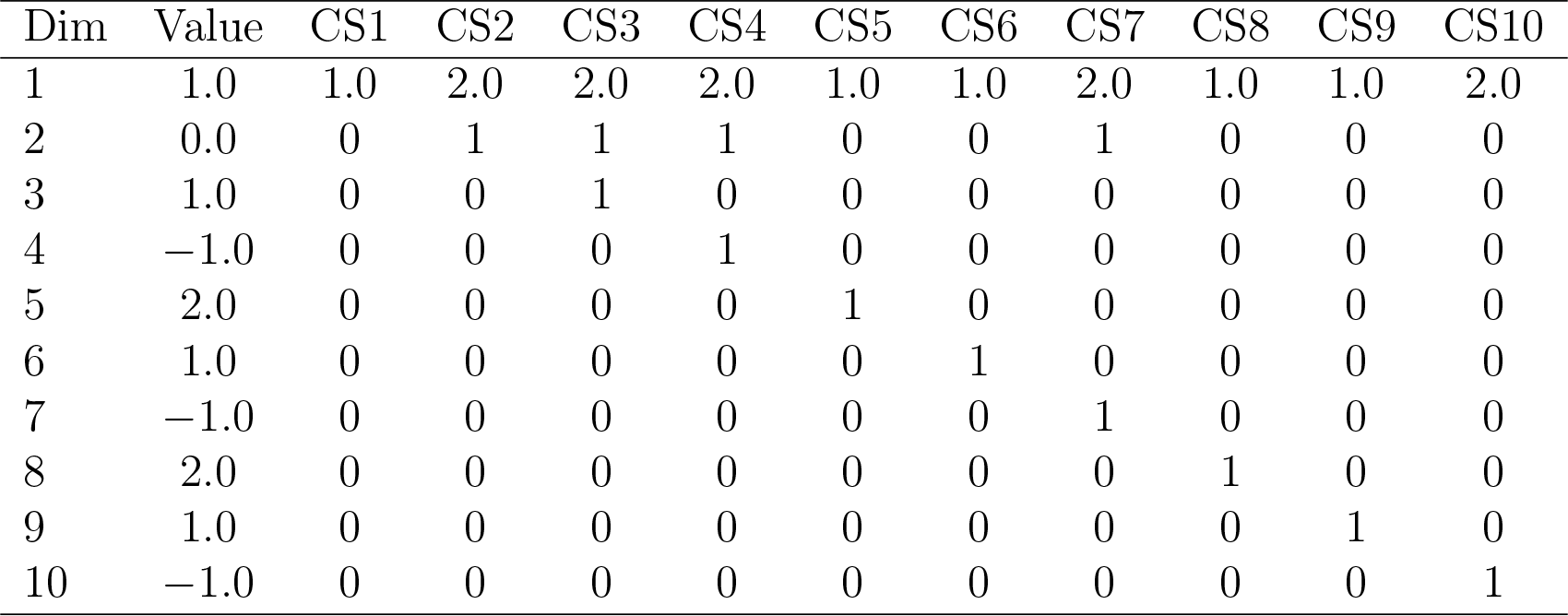
Characteristics of stimulus dimensions and compound stimuli used for the simulation of a change to a more complex world. There are 10 compound stimuli (CS1 to CS10) that can be distinguished using 10 stimulus dimensions. The first dimension represents size, with expected values small (1.0) and large (2.0), and the others are absence/presence (0/1) dimensions. The expected reward values per feature (*W_m_*) are given in the second column and the features of the different compound stimuli are in the following columns.

These compound stimuli, together with six additional compound stimuli, are described in Table 1. Note that we assume log-normal distributions for the first stimulus dimension and also for the stochasticity of rewards, in order to ensure that values are positive.

#### Learning trials

We first consider two cases of sequences of learning trials. In both cases, there is an initial phase of *T* trials of learning (*T* = 1000) with only the first two compound stimuli (e.g., one species of small clients and one species of large clients). This is followed by a phase of an additional *T* trials of learning in a more complex world. In case 1, individuals learn to discriminate between the first four compound stimuli in Table 1 (which could be four client species). In case 2, the world is even more complex, such that individuals learn to discriminate all 10 compound stimuli in Table 1. In both cases, an individual can choose between two compound stimuli in each trial, and these are randomly drawn from all types that occur in that phase of learning of that case.

We also examine a case of reversal learning. In this simulation, there is first a phase of 100 trials where individuals can choose between a rewarded stimulus (*R* = 1), with a feature present in dimension 1, and a non-rewarded stimulus (*R* = 0), with no feature in dimension 1 but a feature present in dimension 2. In practice, the discrimination could be between blue and green stimuli. These 100 trials are enough for individuals to learn to prefer the rewarded stimulus. In the next 100 trials the rewards are reversed. The entire procedure is then repeated for another 200 trials, i.e. an additional two reversals.

Finally, we examine a case of learning set formation. In this simulation, the first 100 trials are as in the reversal learning case, but in subsequent intervals of 100 trials, entirely new pairs of rewarded and non-rewarded stimuli are used, with features in new stimulus dimensions. A total of four pairs are used, making up a total of 400 trials. As an example, the four pairs could be blue and green stimuli, followed by circular and square stimuli, followed by striped and plain stimuli, followed by horizontally and vertically oriented stimuli.

For the above cases, we present results based on replicate simulations of learning for 100 individuals. We assume that the reward from a compound stimulus has a lognormal distribution around the true expected value, with a standard deviation *σ_R_* on the log scale. For the transitions to a more complex world, we use *σ_R_* = 0.10 (in the supplements, we show results for higher stochasticity, *σ_R_* = 0.50; cases 3 to 6), and for reversal learning and learning set formation, we use *σ_R_* = 0.02.

As the starting value of learning rates, we use *α_m_* = 0.04, which allows for learning of unit value differences over 50 to 100 trials. For the starting estimated values, we used *w_m_* = 0; this might hold for individuals without any previous experience of the stimulus dimension.

## Results

### Transitions to a complex world

The first phase of learning for cases 1 and 2, with only two types of compound stimuli, is illustrated in Figure 1. The variation in rewards, shown in Figure 1a, comes both from random variation in the first stimulus dimension (e.g., client size), and from random variation in rewards from a client with a given true expected reward. The sigmoid soft-max curve from equation (4) appears in panel b, and the learning rates *α_m_* and estimated values *w_m_* are shown in panels c and d, averaged over 100 replicates and blocks of 10 learning trials. As seen in Figure 1d, Rescorla-Wagner and Autostep have similar performance in the first phase of learning, with only a slight advantage for Autostep in achieving better estimates of the true values.

There are two cases for the second phase of learning, and the outcome of these learning simulations is illustrated in Figure 2. The learning rates for Autostep increase sharply in the second phase (Figure 2a), especially for case 2 where many new stimulus dimensions are needed for discrimination, whereas the rates for Rescorla-Wagner stay constant. Another comparison of the performance of Rescorla-Wagner and Autostep for the two simulated cases appears in Figure 3. In this figure the performance is measured in terms of the deviation of an individual’s estimate from the true value, implemented as the root mean square error (RMSE). Autostep does noticeably better than Rescorla-Wagner in reducing the errors in the value estimates but, as seen from the Figures 1 and 2, RMSE is not the only thing that matters. Thus, even if a learner deviates in its estimates, it can still be the case that it makes a correct choice between two compound stimuli, because the deviations might be similar for the two stimuli. Quantitatively, over the first 250 post-change trials for case 2, Autostep has a loss of 7.5% of maximum possible reward per trial, whereas Rescorla-Wagner has a higher loss of 11.0%. Over the first 500 post-change trials, these losses are 4.1% and 6.6%. Thus, Autostep does better than Rescorla-Wagner in handling a transition to a more complex world, but the differences are moderate, and not dramatic, seen over timescales of several hundreds of trials.

**Figure 2:**
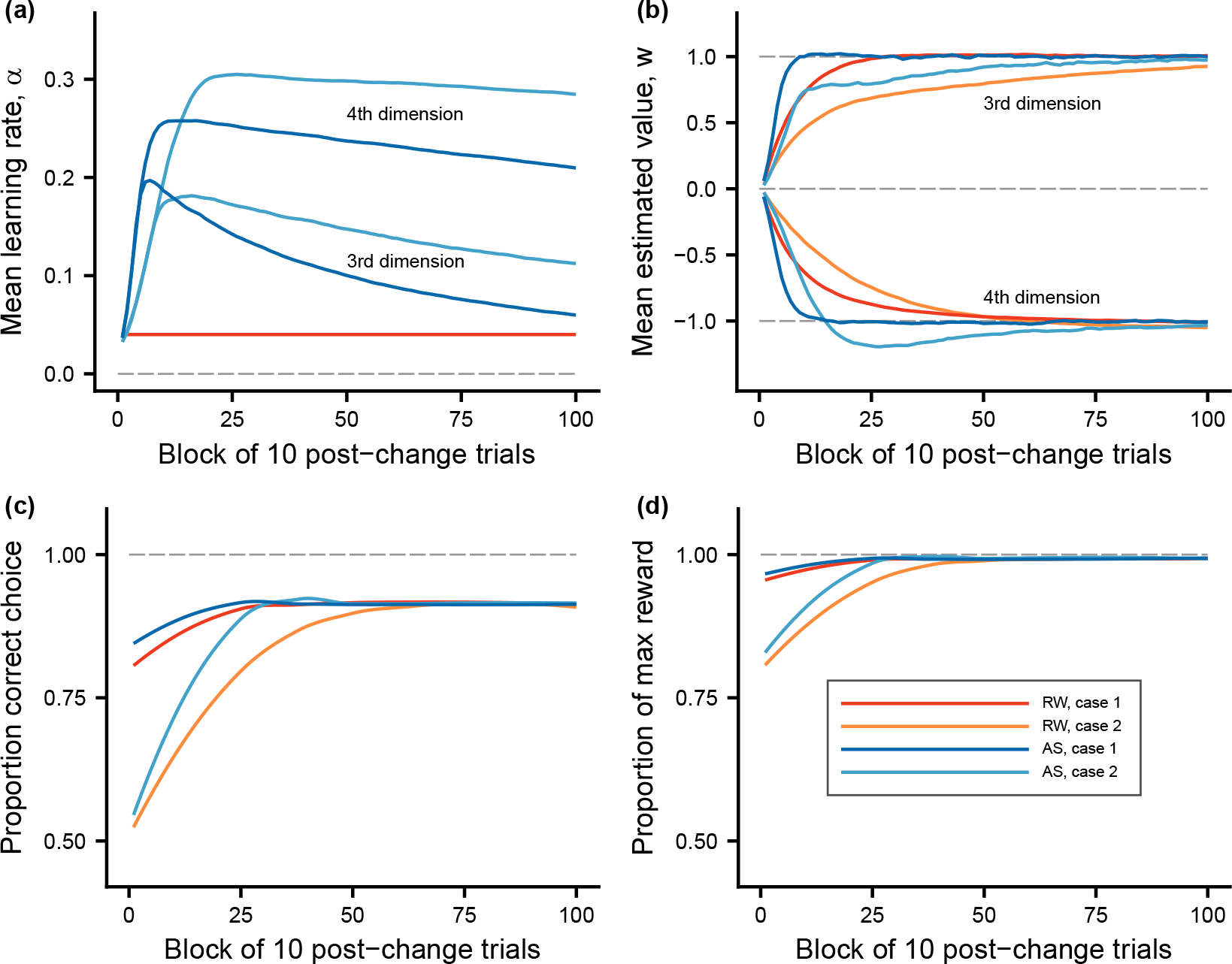
Comparisons of the second phase of learning, when the world becomes more complex, between Rescorla-Wagner (RW) and Autostep (AS), and for the two cases studied. Color coding in panel **(d)** applies to all panels. **(a)** Learning rates for the different learning algorithms and cases (note that the learning rate for Rescorla-Wagner is constant). As an illustration, the third and fourth stimulus dimensions are shown. Note that the features in these dimensions were not present in the first phase. The results are similar for the other new dimensions in Table 1 (dimensions 5-10). **(b)** Estimated values for the different learning algorithms and cases, for stimulus dimensions 3 and 4. **(c)** Proportion of choices that are correct, in the sense of the individual choosing the compound stimulus with higher true value. **(d)** Proportion of reward gained out of the maximum true expected reward available in a trial.

**Figure 3:**
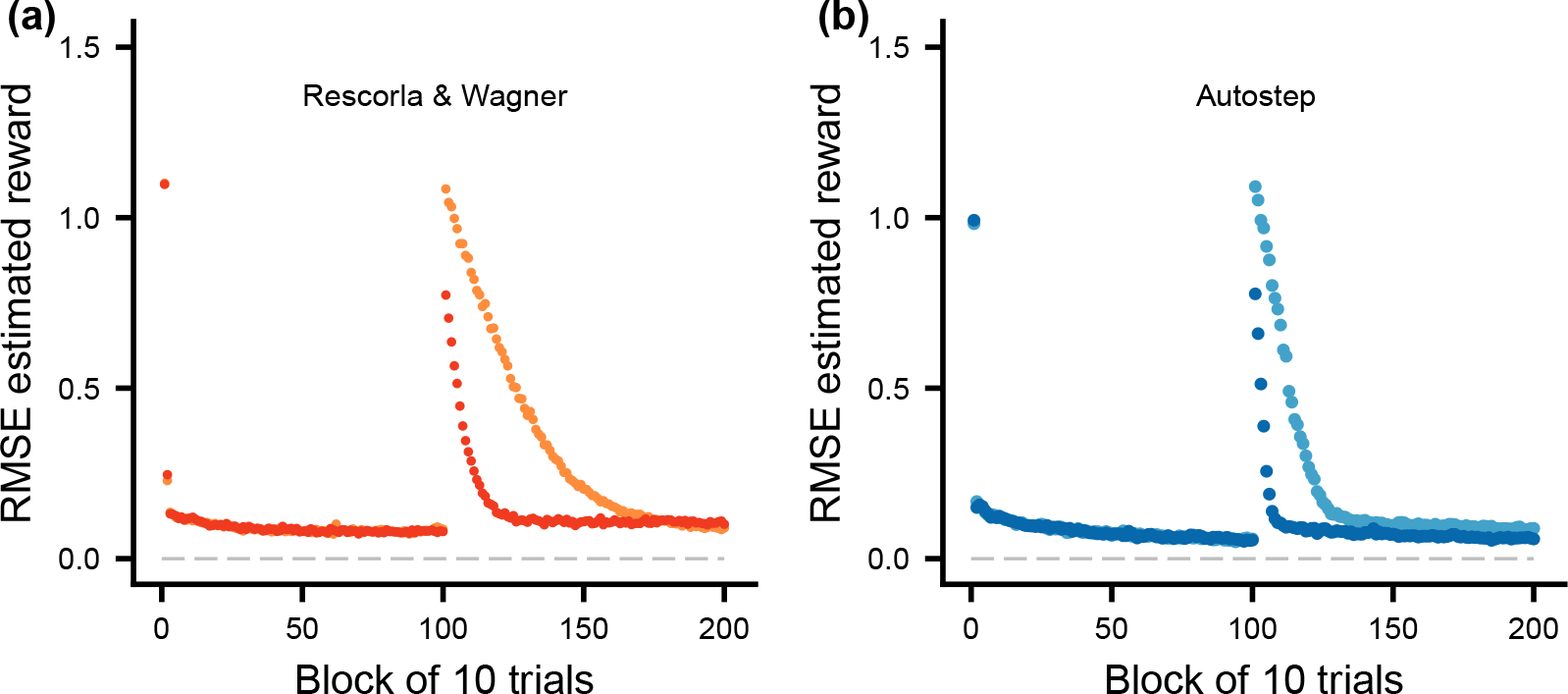
Illustration of the root mean square error (RMSE) of the individual’s estimate (*Q*) of the reward from the selected compound stimulus, plotted against the trial block, over both phases of learning. There are 10 trials in a block and data are averages over 100 replicate learning simulations. (RMSE is similar to a standard deviation but instead measuring the deviation of an estimate from the true value.) **(a)** Rescorla-Wagner learning, with *α*_RW_ = 0.04. **(b)** Autostep learning, following Mahmood et al. (2012). The color coding is as in Figure 2d.

In the supplements we analyze cases similar to those in Figures 1-3, but with high stochasticity (Figures S1-S3). Compared to the cases with lower stochasticity, the learning rates for Autostep are lower, resulting in better estimates of the true values (Figure S3), but there are no dramatic additional advantages for Autostep over Rescorla-Wagner in gaining rewards (Figure S2). Extending the phases of learning to *T* = 10000 trials (Figures S4-S6), the lowering of learning rates for Autostep is even more pronounced, in particular in the first phase of learning (Figure S4). As a result, Autostep does much better than Rescorla-Wagner in estimating the true values (Figure S6).

A different type of analysis of transitions to a more complex world is to consider how much an individual who fails to learn anything new about the more complex world would lose in terms of rewards. For our cases 1 and 2, this would mean that individuals base their choices only on compound stimulus size, also after the transition. Quantitatively, for case 1, where the new world is only moderately more complex, using only size to choose in the second phase would result in a reward loss of around 7.4% per trial, and the corresponding figure for case 2 is around 22% per trial, which is a substantial loss. Note that these losses would apply to all 1000 trials in the second phase, and would be approximately the same for (appropriately modified versions of) Rescorla-Wagner and Autostep. It follows that fairly large advantages can be gained by learning about the new stimulus dimensions in the more complex world.

#### Reversal learning

A comparison of the performance of Rescorla-Wagner and Autostep in reversal learning appears in Fig. 4. For Autostep, the learning rates increase sharply with each successive change in rewards (Figure 4a), whereas for Rescorla-Wagner the learning rates are constant. A consequence of this is that Autostep increases its performance over successive reversals (Figure 4b, c). This means that learning-rate flexibility, for instance as implemented by Autostep, could contribute to observed increases in performance over successive episodes of reversal learning.

**Figure 4:**
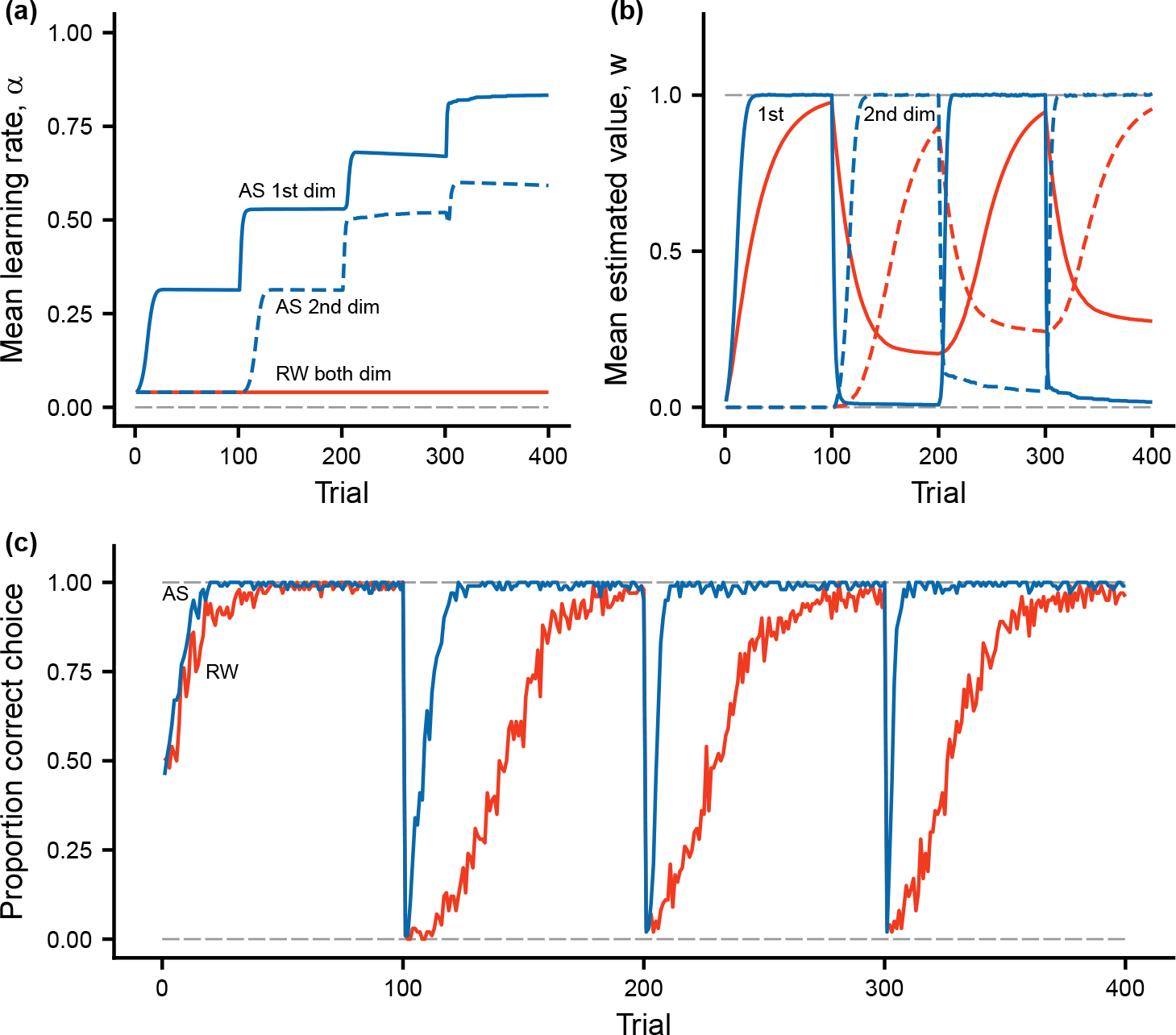
Reversal learning simulation. There are two stimulus dimensions, with 0/1 features that each indicate a type of stimulus. In the first phase of 100 trials, choosing the stimulus with a feature in dimension 1 is rewarded (*R* = 1) and choosing the other, with a feature in dimension 2, is not rewarded (*R* = 0). In the next 100 trials, the rewards are reversed, and then the procedure is repeated for another 200 trials. **(a)** Learning rates for the RescorlaWagner (RW) and Autostep (AS) algorithms (note that the learning rate for Rescorla-Wagner is constant). **(b)** Estimated values for RW and AS, for the two stimulus dimensions. **(c)** Proportion of choices that are correct, in the sense of the individual choosing the stimulus with higher true value.

#### Learning set formation

Figure 5 shows a similar comparison of the performance in learning set formation. In this case there is no additional increase in learning rates for Autostep over successive shifts in pairs of stimuli (Figure 5a), and consequently no increase in performance (Figure 5b, c). Thus, in contrast to reversal learning, learning-rate flexibility does not increase the performance in learning set formation over successive shifts in stimuli to discriminate.

**Figure 5:**
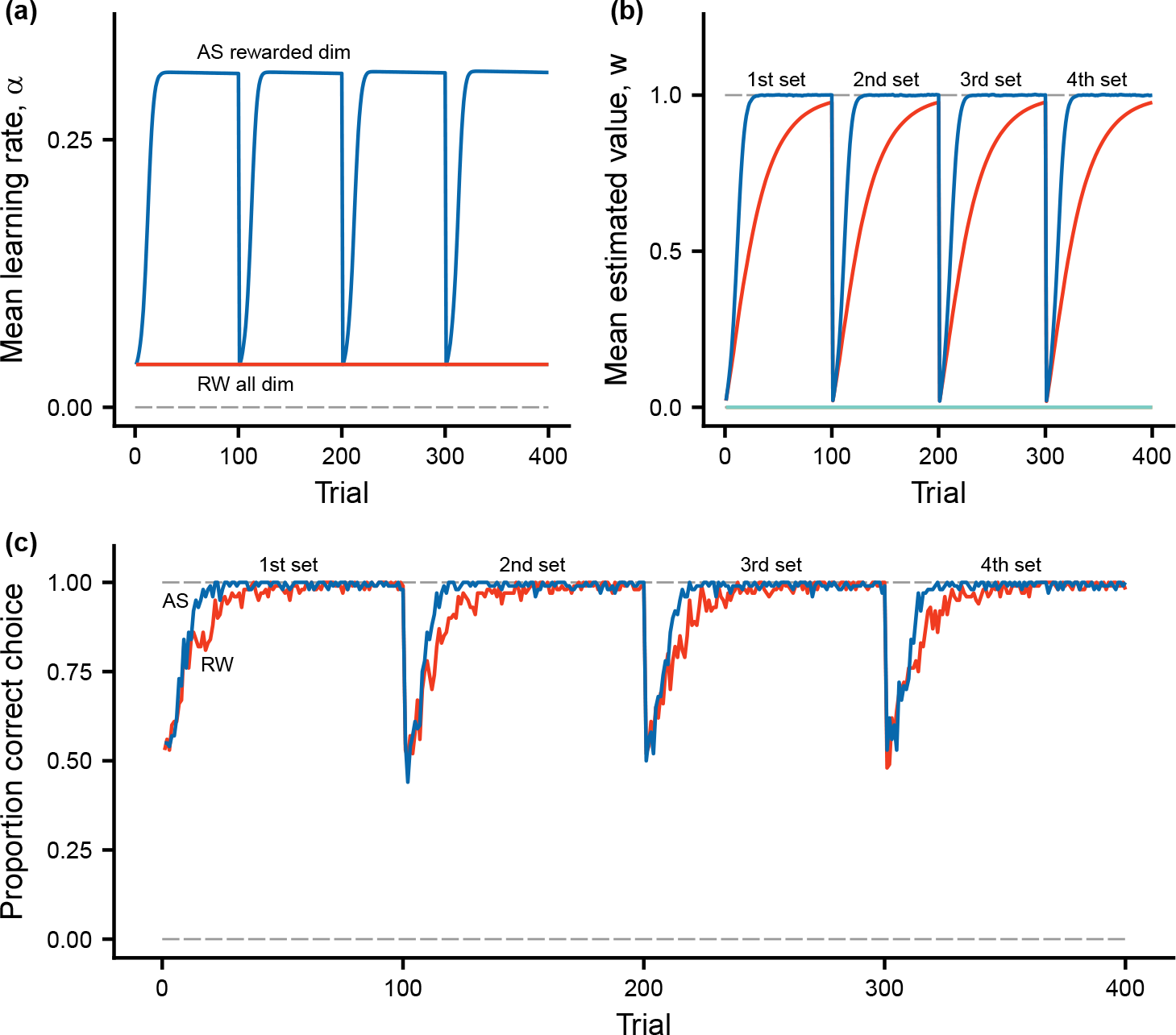
Learning set formation. There are 8 stimulus dimensions, with 0/1 features that each indicate a type of stimulus. In the first phase of 100 trials, there is a choice between the first pair of stimuli, where one is rewarded (*R* = 1) and the other is not rewarded (*R* = 0). In the next 100 trials, a new pair of rewarded and -non-rewarded is used, and so on until four pairs have been used. **(a)** Learning rates for the Rescorla-Wagner (RW) and Autostep (AS) algorithms (note that the learning rate for Rescorla-Wagner is constant). **(b)** Estimated values for RW and AS, for the rewarded stimulus dimension in each set, and for the nonrewarded dimension (green curve at bottom). **(c)** Proportion of choices that are correct, in the sense of the individual choosing the stimulus with higher true value.

## Discussion

In our comparisons of flexible (Autostep) and fixed (Rescorla-Wagner) learning rates, we found pronounced variation in learning rates for the Autostep algorithm (Figures 1c, 2a, 4a, 5a). As a consequence, Autostep performed better than Rescorla-Wagner in estimating the true values of different stimulus dimensions (Figures 1d, 2b, 3, 4b, 5b). For our simulated cases of transitions from a simpler to a more complex world, this meant that Autostep made more correct choices (Figure 2c) and achieved higher post-transition rewards (Figure 2d) than was the case for Rescorla-Wagner. The effects of variable rates on rewards were moderate, but might still be large enough for this kind of learning flexibility to evolve. Alternatively, flexible learning rates could be an aspect of broader cognitive adaptations, relating to attention, memory, and the handling of environmental and social complexity (Emery and Clayton 2004; Deaner et al. 2006; Bond et al. 2007; Izquierdo et al. 2017; Rmus et al. 2021; Leimar et al. 2022).

Our analysis of two frequently used measures of behavioral flexibility produced contrasting results. We found that Autostep increased its performance in reversal learning with each switch in rewards (Figure 4c). This means that a mechanism similar to the one causing the learning rates of Autostep to increase – involving sensitivity to prediction errors that consistently change estimated values over several trials – might contribute to observed improvements in performance in reversal learning with successive switches. In contrast, we found no similar increase in the performance of Autostep over successive shifts of stimuli in learning set formation (Figure 5c). The reason is that, when there is an entirely new situation, in the sense that all stimulus components that a learner encounters come from new stimulus dimensions, the learning rates for Autostep for these dimensions start from scratch. The same ought to hold for other learning mechanisms that do not increase learning rates for stimulus dimensions that a learner so far has not encountered. A tentative conclusion is that increasing performance in reversal learning and in learning set formation correspond to distinct cognitive capacities.

The possibility of different learning rates for different stimulus dimensions is an important aspect of the Rescorla-Wagner model. Its original aim was to explain phenomena such as overshadowing and blocking (Rescorla and Wagner 1972; Miller et al. 1995), and overshadowing of one stimulus component by another depends on differences in learning rates. This is often described in terms of the salience or associability of stimulus components. In nature, the perceived salience of different stimulus components might be adaptive for a particular group of animals. For instance, for some birds the color of artificial prey is more salient than the shape (Kazemi et al. 2014), and such higher learning rates for color might be adaptive. It is learning rate constancy over time, not over stimulus dimensions, that holds for the Rescorla-Wagner model. Our assumption of the same Rescorla-Wagner learning rate for different stimulus dimensions is thus not at all necessary, but is used as a convenient default in the comparison with Autostep.

### Learning models

Many learning models have been proposed in the literature, apart from the ones we study here. Some were developed by experimental psychologists and focus on classical conditioning (e.g., Mackintosh 1975; Pearce and Hall 1980; Le Pelley 2010; Pearce and Mackintosh 2010; Esber and Haselgrove 2011). Although these approaches contain interesting and influential ideas, they turn out not to be suitable for our learning simulations here. The reason is that the specific algorithms have difficulties handling large numbers of stimulus dimension and, furthermore, only allow for fairly limited variation in learning rates.

These approaches discuss variation in learning rates in terms of effects of attention on learning. The idea that attention to stimulus components could be important for learning is often put forward and has been investigated experimentally (Beesley et al. 2015; Niv et al. 2015; Leong et al. 2017; Torrents-Rodas et al. 2021). Nevertheless, models with variation learning rates need not explicitly include attention as a mechanism (Dayan et al. 2000), and Autostep is an example of this.

There are also Kalman-filter-inspired learning models (the Kalman filter originated in the engineering-related fields of optimization and control). Examples are described by Sutton (1992b), Dayan et al. (2000), and Gershman (2015). The Kalman filter gives an optimal solution to a control problem, in certain mathematically well defined situations. It can be used to construct optimal learning algorithms in certain cases where the relative magnitudes of volatility and stochasticity are known (Dayan et al. 2000; Gershman 2015; Piray and Daw 2021). In many situations where the Kalman filter is optimal, the IDBD algorithm achieves approximately the same performance (Sutton 1992b). Because the Autostep algorithm is similar to IDBD, it is reasonable to expect that it has approximately the same performance as a Kalman filter model in situations where the Kalman filter is optimal. A seeming advantage for algorithms such as Autostep and IDBD over a Kalman filter is that they do not require a priori knowledge of the relative magnitudes of volatility and stochasticity.

There is much work in theoretical neuroscience on neural-network-based learning models. This work is of interest if it helps in identifying neural correlates of learning phenomena. An influential example is the modelling by Wang et al. (2018). They present a general perspective on meta learning and report on learning simulations for situations similar to reversal learning and learning set formation. In one simulation they trained a network to obtain rewards in situations with changing volatility, and the network then showed higher learning rates for higher reward volatility, in a similar way as was found in an experiment by Behrens et al. (2007). In another simulation a network was trained on learning set formation, and subsequently showed increasing performance similar to what was found in the original experiments by Harlow (1949). These are interesting results, but it is not clear which kinds of cognitive mechanisms caused the networks to succeed in the learning tasks.

### Behavioral flexibility

The idea that behavioral flexibility should be adaptive in complex worlds is well established. There is evidence that animals that are known or believed to have the cognitive capacities associated with a larger brain, and thus presumably show more flexible behavior, are more successful in novel environments (Sol et al. 2002, 2005). Conversely, there is evidence suggesting that invasive species have cognitive abilities that allow flexible behavior (Szabo et al. 2020). Among the many examples of ecologically relevant situations where there can be a shift to a more complex world are invasions into new habitats (Vardi and Berger-Tal 2022), but also new contextual cues for food choice (Hansen et al. 2010). In nature, individuals are likely to experience environmental changes of many kinds, including the introduction of new significant stimulus dimensions and changes, and even reversals, in the information content of previously encountered stimulus dimensions.

For learning-rate flexibility, which is the focus of our investigation here, it is worth noting that flexibility does not only entail higher learning rates for higher reward volatility, but also lower learning rates for higher reward stochasticity (as seen by comparing Figures 1 and 2 with S1, S2, S4, and S5). While effects of reward stochasticity have been investigated in neuroscience (Nassar et al. 2010; Piray and Daw 2021), there seems to be a lack of studies on the ecological relevance of low vs. high learning rates. Reversal learning experiments, in particular those involving serial reversals, can detect increasing learning rates with repeated reward volatility, as illustrated by our Figure 4. One focus of such studies has been whether species or groups of species differ in this performance. For instance, Bond et al. (2007) compared three corvid species, each of which showed increasing performance with successive reversals, but to different degrees, and suggested that differences in social complexity could explain the observation. Another example is that, based on overviews of several studies, there appears to be a pattern of little or no increase in performance over successive reversals in species of fish (Bitterman 1975; Boussard et al. 2020), in contrast to what is found in other groups of vertebrates. There is so far no well established explanation for this possible difference. In our learning simulations of transitions to a more complex world, we used cleaner fish as an illustrative example, but up to now there are no serial reversal learning experiments on these, and it is not known to what extent they show flexible learning rates. Thus, in principle, cleaner fish learning might be better described by the Rescorla-Wagner model, for instance with adaptive but not very flexible learning rates for different stimulus dimensions, than by the Autostep model. Additional experiments are needed to settle the issue.

A number of studies have examined the neural correlates of learning rate flexibility. One general conclusion is that for humans and non-human primates, as well as for rodents, regions in the prefrontal cortex are important for reversal learning (Izquierdo et al. 2017), with serotonin neurons playing a role (Grossman et al. 2022). In fish, learning experiments on selection lines have shown that brain size influences performance in reversal learning (Buechel et al. 2018) and its decline with age (Boussard et al. 2021), and that specifically relative telencephalon size influences the performance in reversal learning (Triki et al. 2022).

Our simulations showed a qualitative difference between performance in serial reversal learning (Figure 4) and in learning set formation (Figure 5), consistent with the idea that these depend on distinct cognitive capabilities. The capacity for learning set formation appears to have a more narrow phylogenetic distribution, being largely restricted to primates (Harlow 1949; Warren 1966; Deaner et al. 2006) and some species of birds (Wilson et al. 1985; Emery and Clayton 2004; de Mendonça-Furtado and Ottoni 2008), than has increasing performance in serial reversal learning. Learning set formation is sometimes described as rule learning, with the rule being ‘win-stay, lose-shift’ (Warren 1966; Mackintosh et al. 1968; Emery and Clayton 2004), but the actual cognitive mechanism involved is not known. Furthermore, seemingly abstract rules that an experimenter has imposed (e.g., ‘precisely one out of two possibilities is rewarded’) need not correspond to important situations encountered in nature. It might be more important for animals to learn rules about categories of compound stimuli, for instance ‘predator’ and ‘non-predator’. Cleaner fish appear to use this categorization to solve a problem of ‘avoiding punishment’ (Wismer et al. 2016).

Overall, our simulations show that adaptive learning rate flexibility can rely on relatively simple mechanisms, such as using correlations between current and recent changes in estimated values to adjust rates, as for Autostep. To the extent that our examples of transitions to a complex world are biologically realistic, one can also conclude that learning rate flexibility gives a clear but only moderately large advantage over fixed rates. In comparison, as we have shown, it is considerably more important to learn at all about the new and informative stimulus dimensions in the complex world.

Cognitive capacities allowing individuals to achieve this seem essential for behavioral flexibility, and might involve attention, memory, and exploration, in addition to flexible learning rates.

## Acknowledgements

We thank Annika Boussard, Sasha Dall, and Zegni Triki for helpful comments.

## Code availability

C++ source code for the individual-based simulations is available at GitHub, together with instructions for compilation on a Linux operating system: https://github.com/oleimar/learnsim1.

## Supporting information

### Additional figures

**Figure S1:**
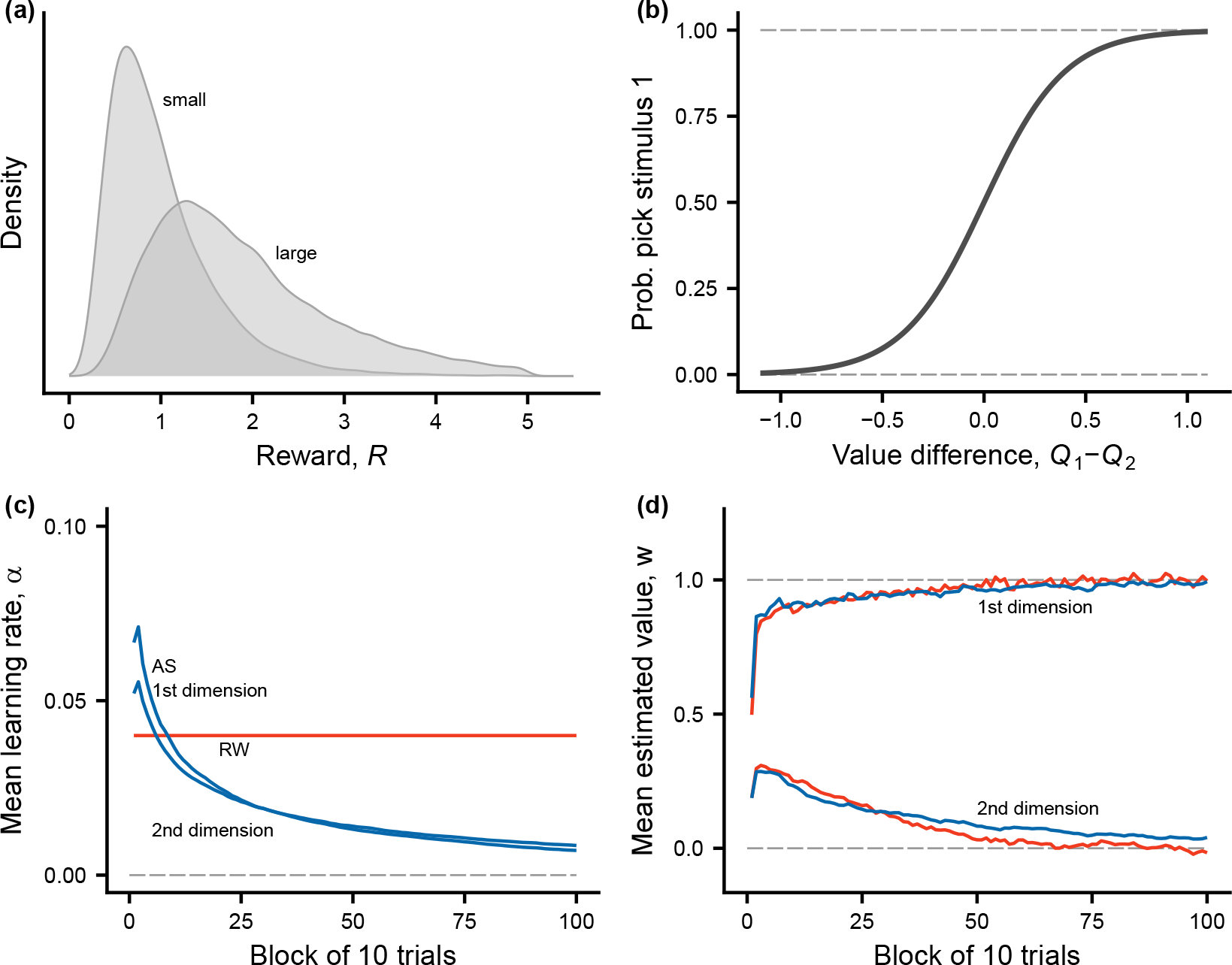
Same as the first cases in the main text (Figure 1), but with high reward stochasticity (*σ_R_* = 0.50). **(a)** Distribution of rewards from the two types of compound stimuli. **(b)** The function from equation (4), giving the probability of choice from the difference in estimated values of the two compound stimuli present in a trial. **(c)** Learning rates for Rescorla-Wagner (RW) and Autostep (AS) for the two stimulus dimensions. **(d)** Estimated values for Rescorla-Wagner and Autostep for the two stimulus dimensions (first dimension has true value 1.0 and second has true value 0). There are 10 trials in a block and data are averages over 100 replicate learning simulations.

**Figure S2:**
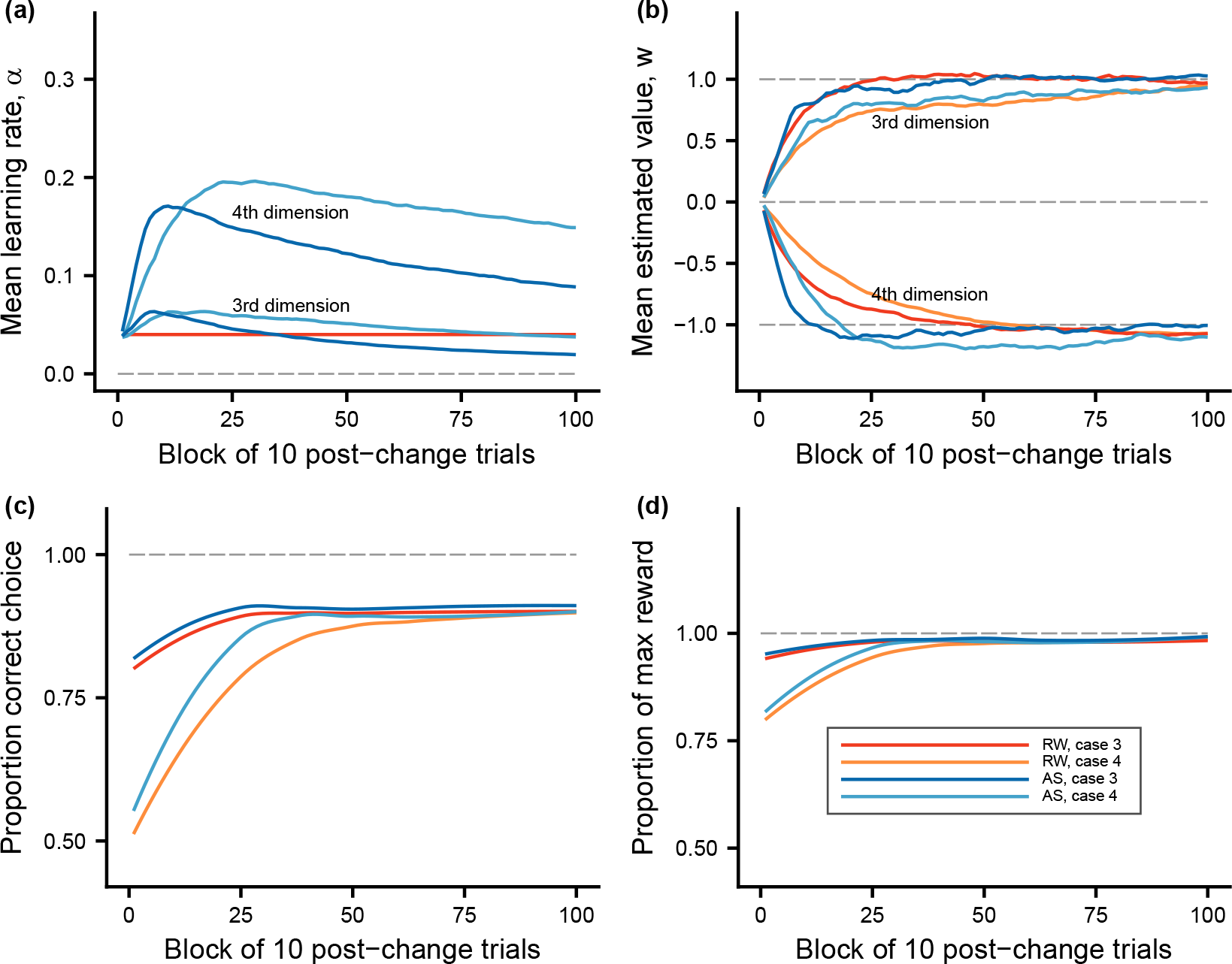
Same as the first cases in the main text (Figure 2), but with high reward stochasticity (*σ_R_*= 0.50). Comparisons of the second phase of learning, when the world becomes more complex, between Rescorla-Wagner (RW) and Autostep (AS), and for the cases studied. Color coding in panel **(d)** applies to all panels. **(a)** Learning rates for the different learning algorithms and cases (note that the learning rate for Rescorla-Wagner is constant). As an illustration, the third and fourth stimulus dimensions are shown. Note that the features in these dimensions were not present in the first phase. **(b)** Estimated values for the different learning algorithms and cases, for stimulus dimensions 3 and 4. **(c)** Proportion of choices that are correct, in the sense of the individual choosing the compound stimulus with higher true value. **(d)** Proportion of reward gained out of the maximum true expected reward available in a trial.

**Figure S3:**
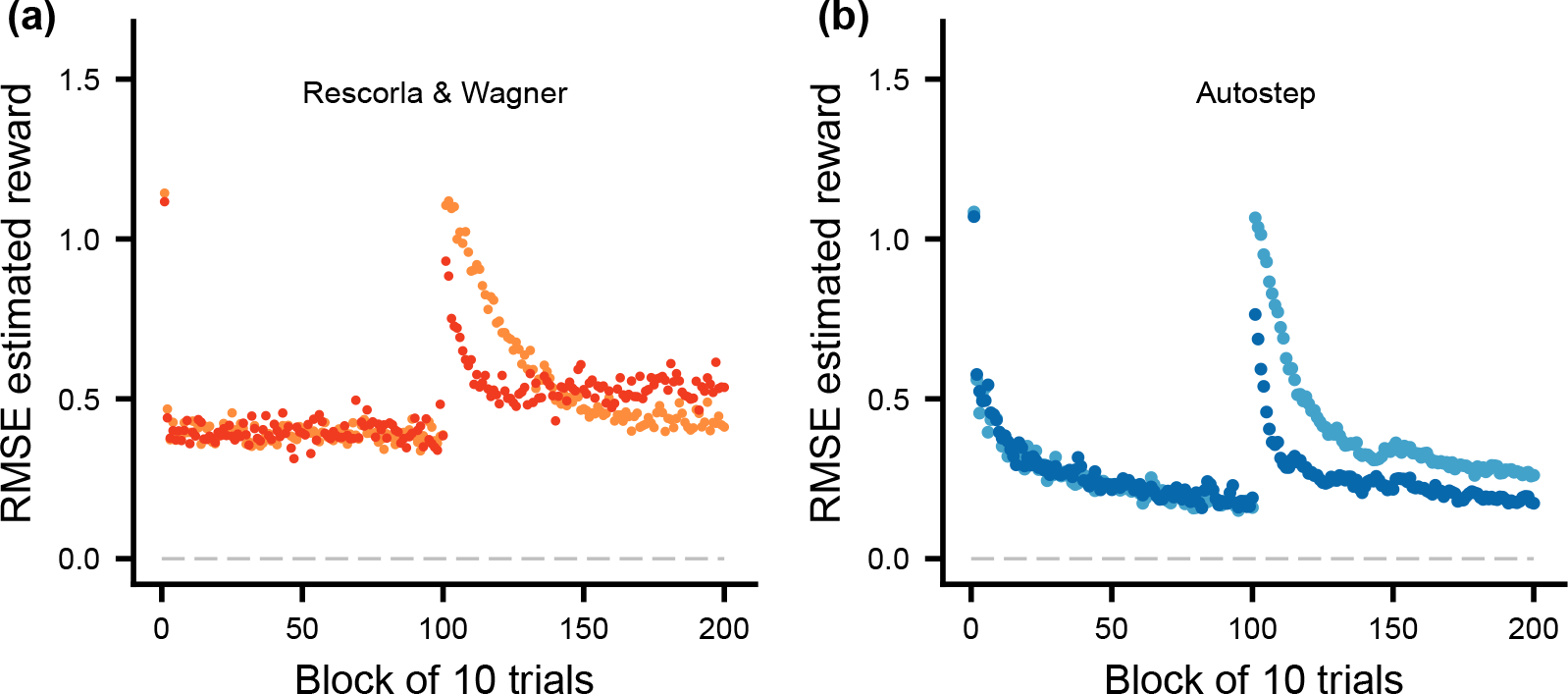
Same as the first cases in the main text (Figure 3), but with high reward stochasticity (*σ_R_*= 0.50). Illustration of the root mean square error (RMSE) of the individual’s estimate (*Q*) of the reward from the selected compound stimulus, plotted against the trial block, over both phases of learning. There are 10 trials in a block and data are averages over 100 replicate learning simulations. **(a)** Rescorla-Wagner learning, with *α*_RW_ = 0.04. **(b)** Autostep learning, following Mahmood et al. (2012). The color coding is as in Figure S2d.

**Figure S4:**
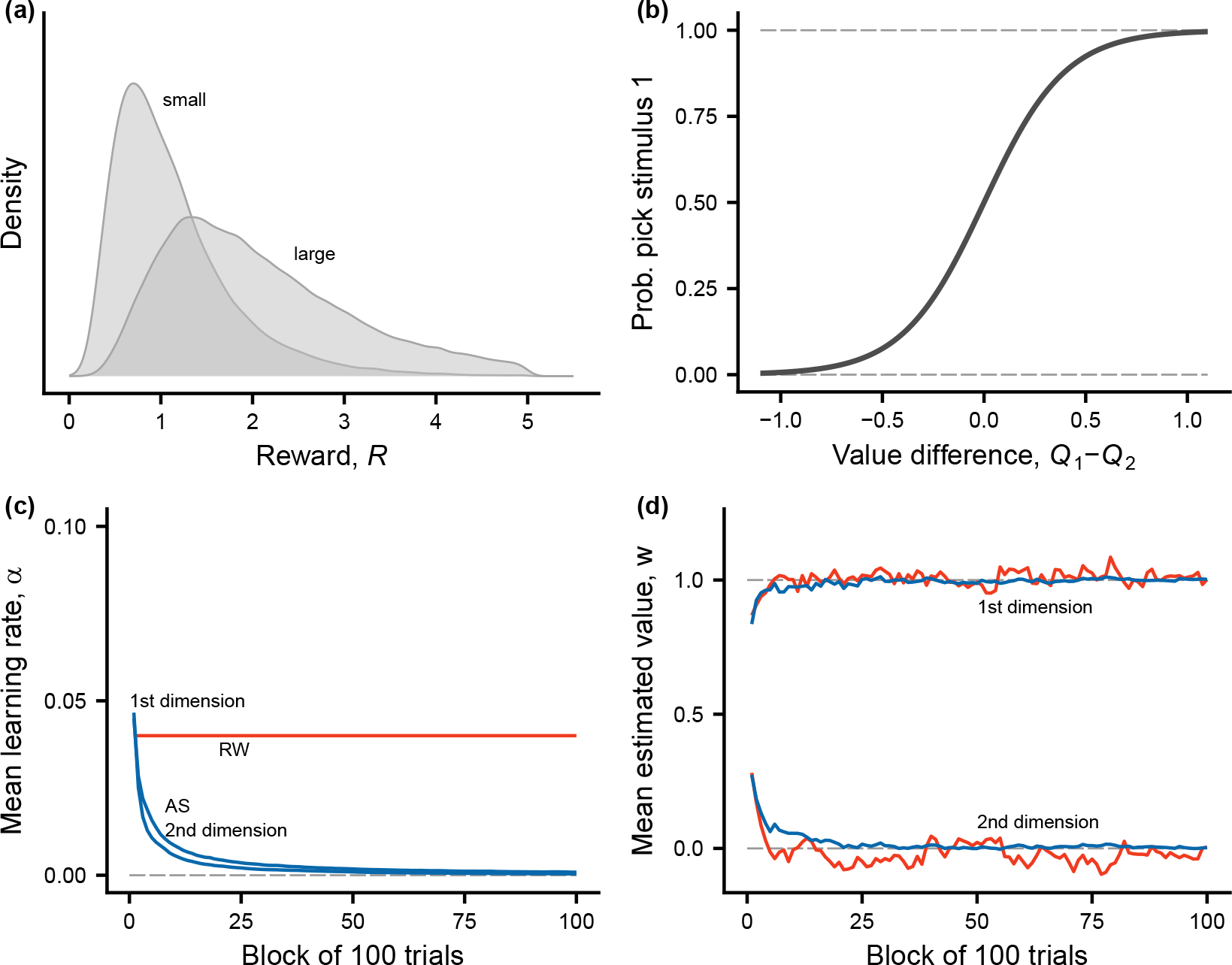
High reward stochasticity (*σ_R_*= 0.50), as in Figure S1, but the first phase of learning is much longer, with *T* = 10000. **(a)** Distribution of rewards from the two types of compound stimuli. **(b)** The function from equation (4), giving the probability of choice from the difference in estimated values of the two compound stimuli present in a trial. **(c)** Learning rates for Rescorla-Wagner (RW) and Autostep (AS) for the two stimulus dimensions. **(d)** Estimated values for Rescorla-Wagner and Autostep for the two stimulus dimensions (first dimension has true value 1.0 and second has true value 0). There are 100 trials in a block and data are averages over 10 replicate learning simulations.

**Figure S5:**
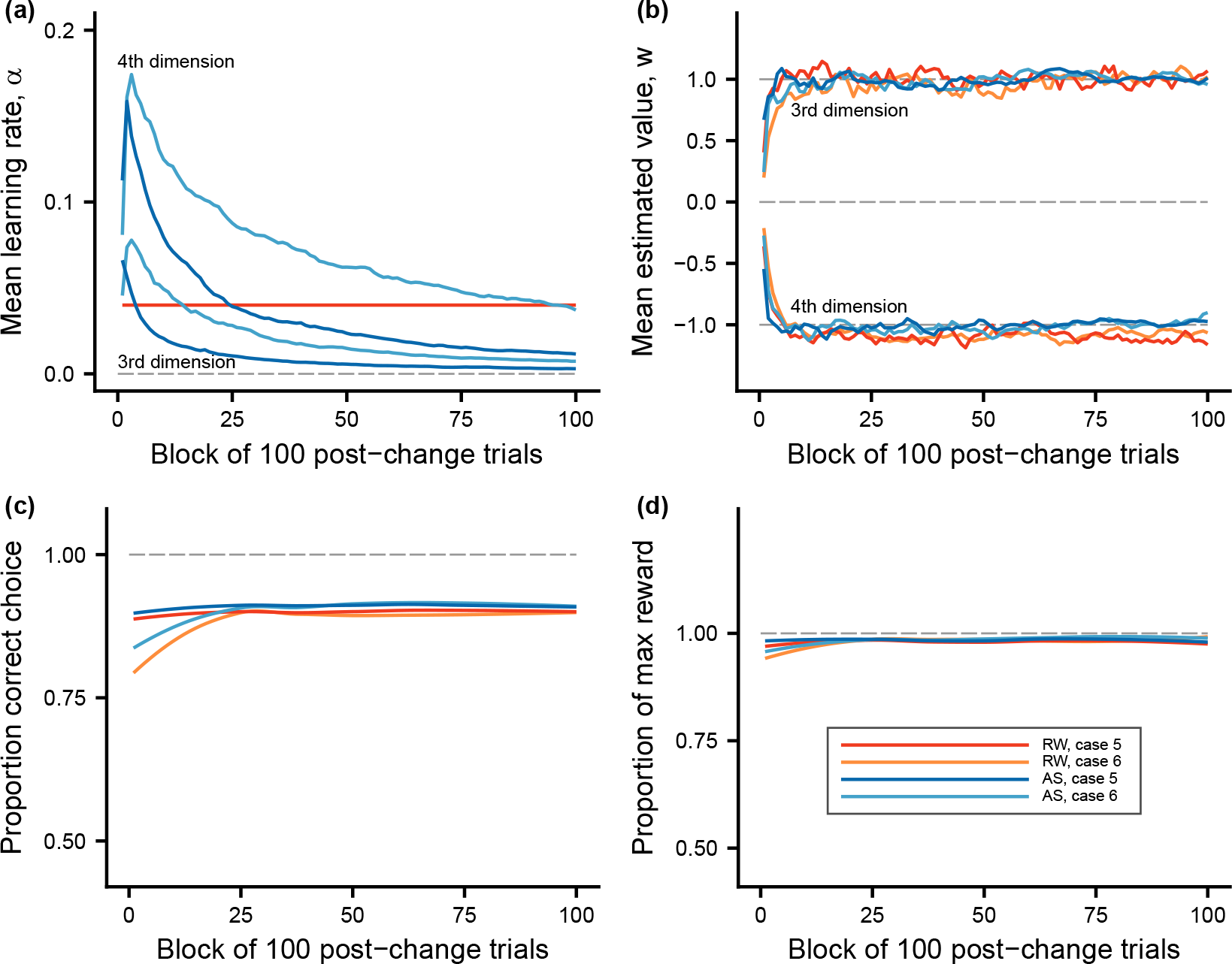
High reward stochasticity (*σ_R_* = 0.50), as in Figure S2, but the second phase of learning is much longer, with *T* = 10000. Comparisons of the second phase of learning, when the world becomes more complex, between Rescorla-Wagner (RW) and Autostep (AS), and for the cases studied. Color coding in panel **(d)** applies to all panels. **(a)** Learning rates for the different learning algorithms and cases (note that the learning rate for Rescorla-Wagner is constant). As an illustration, the third and fourth stimulus dimensions are shown. Note that the features in these dimensions were not present in the first phase. **(b)** Estimated values for the different learning algorithms and cases, for stimulus dimensions 3 and 4. **(c)** Proportion of choices that are correct, in the sense of the individual choosing the compound stimulus with higher true value. **(d)** Proportion of reward gained out of the maximum true expected reward available in a trial.

**Figure S6:**
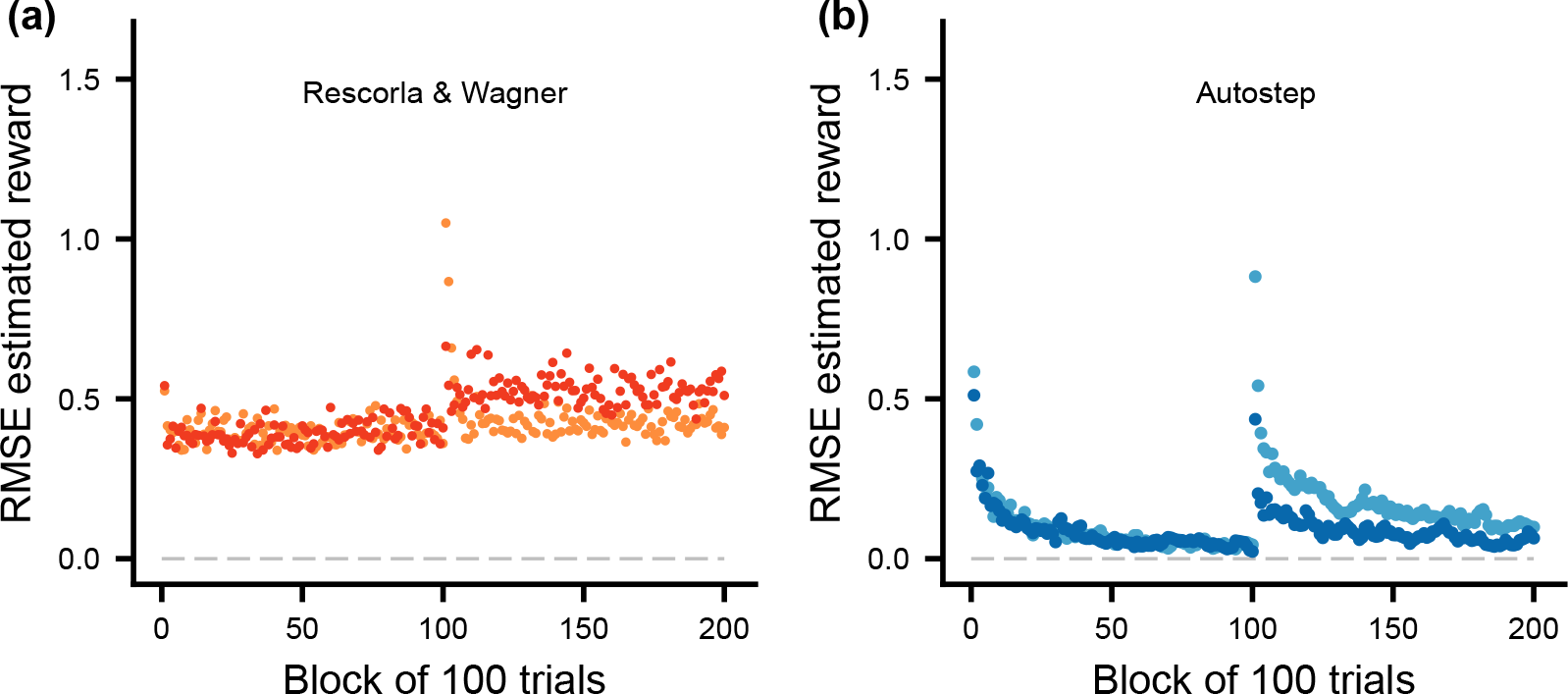
High reward stochasticity (*σ_R_* = 0.50), as in Figure S3, but the first and second phases of learning are much longer, with *T* = 10000. Illustration of the root mean square error (RMSE) of the individual’s estimate (*Q*) of the reward from the selected compound stimulus, plotted against the trial block, over both phases of learning. There are 100 trials in a block and data are averages over 10 replicate learning simulations. **(a)** Rescorla-Wagner learning, with *α*_RW_ = 0.04. **(b)** Autostep learning, following Mahmood et al. (2012). The color coding is as in Figure S5d.

## Model details

Using the notation *W_m_* for the true expected value for a stimulus dimension (e.g., as given in Table 1 of the main text), the true expected reward from a compound stimulus with stimulus components or features *x_m_*, *m* = 1*, . . ., M*, is given by

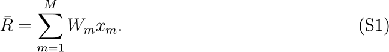

For random variation in the rewards, one possibility is to assume additive variation, but we instead assume that *R* is log-normally distributed, preventing negative values. Thus,

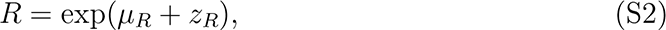

with *µ_R_* such that exp(*µ_R_* + *σ*^2^ */*2) = *R̄*, with *R̄* from equation (S1), i.e. *µ_R_* = log(*R̄*) *− σ*^2^ */*2, with, where *z_R_* is normally distributed with mean zero and standard deviation *σ_R_*. We might, for instance, have *σ_R_* = 0.1, which is used in Figures 1 to 3 in the main text.

Our learning models, as described by equations (1, 2, 3) in the main text and equations (S1, S2) here, are examples of action-value learning. One can view such learning as a modification of classical conditioning, making it applicable to instrumental conditioning (see sections 2.2 and 2.5 in Sutton and Barto (2018) for discussion of this learning approach). Note also that action-value learning can be regarded as a simplified version of the Sarsa algorithm, for cases where individuals do not use any sophisticated states and where each learning trial is a separate episode (terminology from Sutton and Barto (2018)). We get a connection to the presentation in Sutton and Barto (2018) by assuming that the state in a trial is just the compound stimuli that are present in that trial, for the individual for choose between.

## Learning rates

### Rescorla-Wagner

The Rescorla-Wagner learning mechanism has learning rates that are constant in time.

In our simulations, we assume that

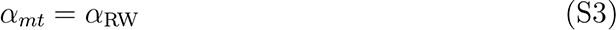

for the learning rate for dimension *m* in trial *t*. Thus, we assume that the RescorlaWagner learning rate *α*_RW_ is constant, and thus independent of the stimulus dimension *m* and the trial *t* (we use *α*_RW_ = 0.04 in our simulations).

### IDBD

The IDBD (Incremental Delta-Bar-Delta) learning algorithm was developed by Sutton (1992a), and is also described in Sutton (1992b) and by equations (1) and (3) in Mahmood et al. (2012). For clarity we write the prediction error from equation (2) in trial *t* as

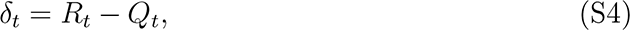

where *Q_t_* is the estimated value from equation (3) for the chosen compound stimulus with stimulus components *x_m_*. We write the estimated value as

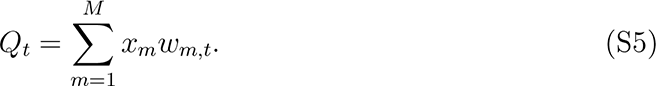

Using the order of updates for IDBD from Sutton (1992a,b) and Mahmood et al. (2012), there are initial learning rates *α_m,_*_1_ and, to update the estimated values, we first update the learning rates as follows:

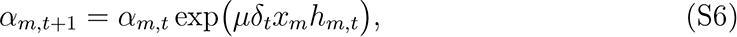

where *µ* is a meta learning rate, *δ_t_*is from equation (S4), and *h_m,t_*is an additional quantity with starting value *h_m,_*_1_ = 0. The estimated value updates, corresponding to equation (1), are then

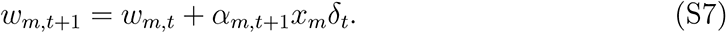

The quantity *h_m,t_*is updated as follows

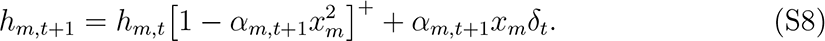

The notation [X]^+^ means equal to *X* for positive *X* and zero otherwise.

For an intuitive interpretation, note the quantity *h_m_* is a kind of memory or trace of changes to *w_m_*, because those changes are given by *α_m,t_*_+1_*x_m_δ_t_*. The exponent in equation (S6) contains the product of this trace with *δ_t_x_m_*, which is proportional to the current change in *w_m_*. Thus, if successive changes to *w_m_*tend to be positively correlated, the learning rate *α_m_* will increase, and similarly decrease for negative correlations.

### Autostep

A problem with the IDBD algorithm, pointed out by Mahmood et al. (2012), is that it is very sensitive to the exact value of the meta learning rate *µ*. To avoid this problem, Mahmood et al. (2012) introduced changes and obtained a more robust algorithm, which they called Autostep. It is called Autostep because the step size, or effective meta-learning rate, is automatically adjusted to an appropriate value. The Autostep algorithm is given in Table 1 of Mahmood et al. (2012), and is as follows.

First set the two meta learning parameters *µ* and *τ* (we used *µ* = 0.2 and *τ* = 100 in our Autostep simulations). Then initialise *w_m,_*_1_ and *α_m,_*_1_ (we used *w_m,_*_1_ = 0 and *α_m,_*_1_ = 0.04) and set *v_m,_*_1_ = 0 and *h_m,_*_1_ = 0.

For each trial *t* we then have the following. The estimated values for compound stimuli are computed from features *x_m_* as

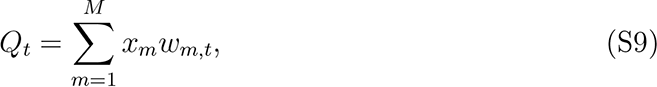

just as previously. The learner chooses a compound stimulus using the soft-max rule in equation (4) and perceives a reward *R*. This gives a prediction error

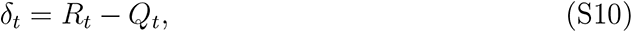

just as previously. Then compute the quantity

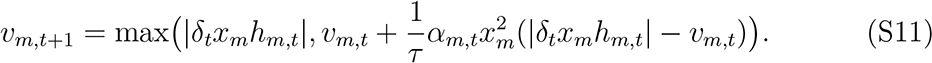

If *v_m,t_*_+1_ */*= 0, compute

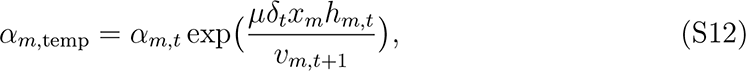

otherwise use *α_m,_*_temp_ = *α_m,t_*. Do this for each *m* and compute

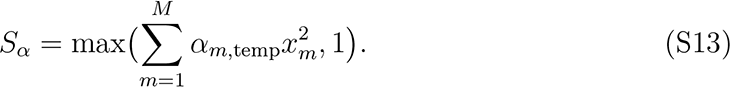

Use this *S_α_*to normalise the learning rates

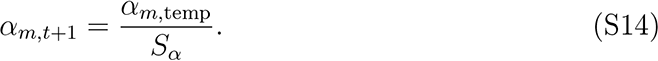

These are then used for the updates of the estimated values:

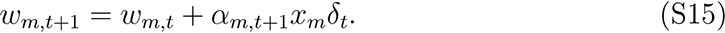

Finally, the *h_m,t_*are updated as follows:

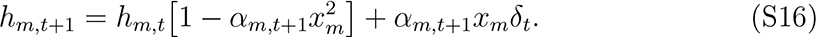

The changes in Autostep from IDBD are, first, that the exponent in the update in equation (S6) is divided by the quantity *v_m,t_*_+1_ in equation (S12), achieving a reasonable effective meta step size and, second, that the learning rates are normalised in equation (S14). See Mahmood et al. (2012) for more comments on the Autostep algorithm.

## References

1. Audet, J.-N. and Lefebvre, L. 2017. What’s flexible in behavioral flexibility? Behavioral Ecology, 28(4):943–947.

2. Bailey, A. M., McDaniel, W. F., and Thomas, R. K. 2007. Approaches to the study of higher cognitive functions related to creativity in nonhuman animals. Methods, 42(1):3–11.

3. Bairlein, F. and Simons, D. 1995. Nutritional adaptations in migrating birds. Israel Journal of Ecology and Evolution, 41(3):357–367.

4. Beesley, T., Nguyen, K. P., Pearson, D., and Le Pelley, M. E. 2015. Uncertainty and predictiveness determine attention to cues during human associative learning. Quarterly Journal of Experimental Psychology, 68(11):2175–2199.

5. Behrens, T. E. J., Woolrich, M. W., Walton, M. E., and Rushworth, M. F. S. 2007. Learning the value of information in an uncertain world. Nature Neuroscience, 10(9):1214–1221.

6. Bitterman, M. E. 1975. The comparative analysis of learning. Science, 188(4189):699– 709.

7. Bond, A. B., Kamil, A. C., and Balda, R. P. 2007. Serial reversal learning and the evolution of behavioral flexibility in three species of North American corvids (Gymnorhinus cyanocephalus, Nucifraga columbiana, Aphelocoma californica). Journal of Comparative Psychology, 121(4):372–379.

8. Boussard, A., Amcoff, M., Buechel, S. D., Kotrschal, A., and Kolm, N. 2021. The link between relative brain size and cognitive ageing in female guppies (*Poecilia reticulata*) artificially selected for variation in brain size. Experimental Gerontology, 146:111218.

9. Boussard, A., Buechel, S. D., Amcoff, M., Kotrschal, A., and Kolm, N. 2020. Brain size does not predict learning strategies in a serial reversal learning test. Journal of Experimental Biology, 223(15):jeb224741.

10. Buechel, S. D., Boussard, A., Kotrschal, A., van der Bijl, W., and Kolm, N. 2018. Brain size affects performance in a reversal-learning test. Proceedings of the Royal Society B: Biological Sciences, 285(1871):20172031.

11. Cauchoix, M., Hermer, E., Chaine, A. S., and Morand-Ferron, J. 2017. Cognition in the field: comparison of reversal learning performance in captive and wild passerines. Scientific Reports, 7(1):12945.

12. Dayan, P., Kakade, S., and Montague, P. R. 2000. Learning and selective attention. Nature Neuroscience, 3(11):1218–1223.

13. de Mendonça-Furtado, O. and Ottoni, E. B. 2008. Learning generalization in problem solving by a blue-fronted parrot (Amazona aestiva). Animal Cognition, 11(4):719– 725.

14. Deaner, R. O., van Schaik, C. P., and Johnson, V. 2006. Do some taxa have better domain-general cognition than others? A meta-analysis of nonhuman primate studies. Evolutionary Psychology, 4(1):147470490600400114.

15. Diederen, K. M. J. and Schultz, W. 2015. Scaling prediction errors to reward variability benefits error-driven learning in humans. Journal of Neurophysiology, 114(3):1628– 1640.

16. Emery, N. J. and Clayton, N. S. 2004. The mentality of crows: convergent evolution of intelligence in corvids and apes. Science, 306(5703):1903–1907.

17. Esber, G. R. and Haselgrove, M. 2011. Reconciling the influence of predictiveness and uncertainty on stimulus salience: a model of attention in associative learning. Proceedings of the Royal Society B: Biological Sciences, 278(1718):2553–2561.

18. Farashahi, S., Xu, J., Wu, S.-W., and Soltani, A. 2020. Learning arbitrary stimulusreward associations for naturalistic stimuli involves transition from learning about features to learning about objects. Cognition, 205:104425.

19. Fawcett, T. W., Hamblin, S., and Giraldeau, L. A. 2013. Exposing the behavioral gambit: The evolution of learning and decision rules. Behavioral Ecology, 24(1):2– 11.

20. Gershman, S. J. 2015. A unifying probabilistic view of associative learning. PLOS Computational Biology, 11(11):e1004567.

21. Grossman, C. D., Bari, B. A., and Cohen, J. Y. 2022. Serotonin neurons modulate learning rate through uncertainty. Current Biology, 32(3):586–599.e7.

22. Grutter, A. S. 1994. Spatial and temporal variations of the ectoparasites of seven reef fish species from lizard island and heron island, australia. Marine Ecology Progress Series, 115:21–30.

23. Grutter, A. S. 1995. Relationship between cleaning rates and ectoparasite loads in coral reef fishes. Marine Ecology Progress Series, 118:51–58.

24. Grutter, A. S. and Bshary, R. 2004. Cleaner fish, *Labroides dimidiatus*, diet preferences for different types of mucus and parasitic gnathiid isopods. Animal Behaviour, 68(3):583–588.

25. Hansen, B. T., Holen, O. H., and Mappes, J. 2010. Predators use environmental cues to discriminate between prey. Behavioral Ecology and Sociobiology, 64(12):1991–1997.

26. Harlow, H. F. 1949. The formation of learning sets. Psychological Review, 56(1):51–65.

27. Holland, P. C. and Schiffino, F. L. 2016. Mini-review: Prediction errors, attention and associative learning. Neurobiology of Learning and Memory, 131:207–215.

28. Izquierdo, A., Brigman, J. L., Radke, A. K., Rudebeck, P. H., and Holmes, A. 2017. The neural basis of reversal learning: An updated perspective. Neuroscience, 345:12–26.

29. Jacobs, R. A. 1988. Increased rates of convergence through learning rate adaptation. Neural Networks, 1(4):295–307.

30. Janmaat, K. R., Boesch, C., Byrne, R., Chapman, C. A., Goné Bi, Z. B., Head, J. S., Robbins, M. M., Wrangham, R. W., and Polansky, L. 2016. Spatio-temporal complexity of chimpanzee food: How cognitive adaptations can counteract the ephemeral nature of ripe fruit. American Journal of Primatology, 78(6):626–645.

31. Kazemi, B., Gamberale-Stille, G., Tullberg, B. S., and Leimar, O. 2014. Stimulus salience as an explanation for imperfect mimicry. Current Biology, 24(9):965–969.

32. Le Pelley, M. E. 2010. The hybrid modeling approach to conditioning. In Schmajuk, N., editor, Computational Models of Conditioning, pages 71–107. Cambridge University Press, Cambridge, UK.

33. Lea, S. E. G., Chow, P. K. Y., Leaver, L. A., and McLaren, I. P. L. 2020. Behavioral flexibility: A review, a model, and some exploratory tests. Learning & Behavior, 48(1):173–187.

34. Leimar, O., Dall, S. R. X., Houston, A. I., and McNamara, J. M. 2022. Behavioural specialization and learning in social networks. Proceedings of the Royal Society B: Biological Sciences, 289(1980):20220954.

35. Leong, Y. C., Radulescu, A., Daniel, R., DeWoskin, V., and Niv, Y. 2017. Dynamic interaction between reinforcement learning and attention in multidimensional environments. Neuron, 93(2):451–463.

36. Liu, Y., Day, L. B., Summers, K., and Burmeister, S. S. 2016. Learning to learn: advanced behavioural flexibility in a poison frog. Animal Behaviour, 111:167–172.

37. Mackintosh, N. J. 1975. A theory of attention: Variations in the associability of stimuli with reinforcement. Psychological Review, 82(4):276–298.

38. Mackintosh, N. J., Mcgonigle, B., and Holgate, V. 1968. Factors underlying improvement in serial reversal learning. Canadian Journal of Psychology, 22(2):85–95.

39. Mahmood, A. R., Sutton, R. S., Degris, T., and Pilarski, P. M. 2012. Tuning-free step-size adaptation. In 2012 IEEE International Conference on Acoustics, Speech and Signal Processing (ICASSP), pages 2121–2124.

40. Miller, R. R., Barnet, R. C., and Grahame, N. J. 1995. Assessment of the RescorlaWagner model. Psychological Bulletin, 117(3):363–386.

41. Nassar, M. R., Wilson, R. C., Heasly, B., and Gold, J. I. 2010. An approximately Bayesian delta-rule model explains the dynamics of belief updating in a changing environment. Journal of Neuroscience, 30(37):12366–12378.

42. Niv, Y., Daniel, R., Geana, A., Gershman, S. J., Leong, Y. C., Radulescu, A., and Wilson, R. C. 2015. Reinforcement learning in multidimensional environments relies on attention mechanisms. The Journal of Neuroscience, 35(21):8145–8157.

43. Pearce, J. M. and Hall, G. 1980. A model for Pavlovian learning: Variations in the effectiveness of conditioned but not of unconditioned stimuli. Psychological Review, 87(6):532–552.

44. Pearce, J. M. and Mackintosh, N. J. 2010. Two theories of attention: a review and possible integration. In Mitchell, C. J. and Le Pelley, M. E., editors, Attention and Associative Learning: From Brain to Behaviour, pages 11–40. Oxford University Press, Oxford, UK.

45. Pierce, B. J. and McWilliams, S. R. 2005. Seasonal changes in composition of lipid stores in migratory birds: causes and consequences. The Condor, 107(2):269–279.

46. Piray, P. and Daw, N. D. 2021. A model for learning based on the joint estimation of stochasticity and volatility. Nature Communications, 12(1):6587.

47. Raine, N. E. and Chittka, L. 2012. No trade-off between learning speed and associative flexibility in bumblebees: a reversal learning test with multiple colonies. PLOS ONE, 7(9):e45096.

48. Rescorla, R. A. and Wagner, A. R. 1972. A theory of Pavlovian conditioning: Variations in the effectiveness of reinforcement and nonreinforcement. In Black, A. H. and Prokasy, W. F., editors, Classical conditioning II: current research and theory, pages 64–99. Appleton-Century-Crofts, New York.

49. Rmus, M., McDougle, S. D., and Collins, A. G. 2021. The role of executive function in shaping reinforcement learning. Current Opinion in Behavioral Sciences, 38:66–73.

50. Roche, D. G., Jornod, M., Douet, V., Grutter, A. S., and Bshary, R. 2021. Client fish traits underlying variation in service quality in a marine cleaning mutualism. Animal Behaviour, 175:137–151.

51. Sol, D., Duncan, R. P., Blackburn, T. M., Cassey, P., and Lefebvre, L. 2005. Big brains, enhanced cognition, and response of birds to novel environments. Proceedings of the National Academy of Sciences, 102(15):5460–5465.

52. Sol, D., Timmermans, S., and Lefebvre, L. 2002. Behavioural flexibility and invasion success in birds. Animal Behaviour, 63(3):495–502.

53. Soltani, A. and Izquierdo, A. 2019. Adaptive learning under expected and unexpected uncertainty. Nature Reviews Neuroscience, 20(10):635–644.

54. Sutton, R. S. 1992a. Adapting bias by gradient descent: An incremental version of delta-bar-delta. In Proceedings of the Tenth National Conference on Artificial Intelligence, pages 171–176. MIT Press, Cambridge, MA.

55. Sutton, R. S. 1992b. Gain adaptation beats least squares? In Proceedings of the Seventh Yale Workshop on Adaptive and Learning Systems, pages 161–166. Yale University, New Haven, CT.

56. Sutton, R. S. 2022. A history of meta-gradient: Gradient methods for meta-learning. arXiv preprint, 2202.09701, pages 1–8.

57. Sutton, R. S. and Barto, A. G. 2018. Reinforcement learning: An introduction second edition. MIT Press, Cambridge, MA.

58. Szabo, B., Damas-Moreira, I., and Whiting, M. J. 2020. Can cognitive ability give invasive species the means to succeed? A review of the evidence. Frontiers in Ecology and Evolution, 8:187.

59. Torrents-Rodas, D., Koenig, S., Uengoer, M., and Lachnit, H. 2021. Evidence for two attentional mechanisms during learning. Quarterly Journal of Experimental Psychology, 74(12):2112–2123.

60. Triki, Z., Granell-Ruiz, M., Fong, S., Amcoff, M., and Kolm, N. 2022. Brain morphology correlates of learning and cognitive flexibility in a fish species (*Poecilia reticulata)*. Proceedings of the Royal Society B: Biological Sciences, 289(1978):20220844.

61. Triki, Z., Wismer, S., Rey, O., Ann Binning, S., Levorato, E., and Bshary, R. 2019. Biological market effects predict cleaner fish strategic sophistication. Behavioral Ecology, 30(6):1548–1557.

62. Trimmer, P. C., McNamara, J. M., Houston, A. I., and Marshall, J. A. R. 2012. Does natural selection favour the Rescorla–Wagner rule? Journal of Theoretical Biology, 302:39–52.

63. Uddin, L. Q. 2021. Cognitive and behavioural flexibility: neural mechanisms and clinical considerations. Nature Reviews Neuroscience, 22(3):167–179.

64. Vardi, R. and Berger-Tal, O. 2022. Environmental variability as a predictor of behavioral flexibility in urban environments. Behavioral Ecology, 33(3):573–581.

65. Wang, J. X., Kurth-Nelson, Z., Kumaran, D., Tirumala, D., Soyer, H., Leibo, J. Z., Hassabis, D., and Botvinick, M. 2018. Prefrontal cortex as a meta-reinforcement learning system. Nature Neuroscience, 21(6):860–868.

66. Warren, J. M. 1966. Reversal learning and the formation of learning sets by cats and rhesus monkeys. Journal of Comparative and Physiological Psychology, 61(3):421– 428.

67. Wilson, B., Mackintosh, N. J., and Boakes, R. A. 1985. Transfer of relational rules in matching and oddity learning by pigeons and corvids. The Quarterly Journal of Experimental Psychology Section B, 37(4b):313–332.

68. Wismer, S., Grutter, A., and Bshary, R. 2016. Generalized rule application in bluestreak cleaner wrasse (Labroides dimidiatus): using predator species as social tools to reduce punishment. Animal Cognition, 19(4):769–778.

